# Antifungal activity of the novel compound M451 against phytopathogens

**DOI:** 10.1101/2023.02.04.525039

**Authors:** V Tetz, K Kardava, K Krasnov, M Vecherkovskaya, G Tetz

## Abstract

Phytopathogenic fungi are the dominant causal agents of plant diseases. Currently available fungicides have significant disadvantages, being insufficiently effective owing to both intrinsic tolerance and the spread of antibiotic resistance accumulating in plant tissues, posing a global threat to public health. Finding a new broad-spectrum fungicide is a challenge for plant protection. We studied the potency of a novel antimicrobial agent, M451, against different phytopathogenic fungi of the phyla *Ascomycota, Oomycota*, and *Basidiomycota*. M451 exhibited significant antifungal activity with EC50 values ranging from 34 to 145 µg/mL. Analysis of the minimal fungicidal concentration and conidial destruction assay revealed that M451 possesses the highest activity compared with different polyene, azole, and phenylpyrrole antifungals against *Fusarium oxysporum*. Time-kill analysis revealed that M451 was the only antifungal agent tested that exhibited antifungal activity within 5 min of exposure. Spore production and germination were also significantly inhibited by M451 treatment. Based on the broad spectrum of antifungal effects across different plant pathogens, M451 could be a new chemical fungicide for plant disease management.

## 1 Introduction

Phytopathogenic fungi are among the dominant causal agents of plant diseases, and can seriously interrupt the normal growth of crops, fruits, and vegetables (Fisher M.C. et al., 2012; Doehlemann G. et al., 2017; Jain A. et al., 2019; Blackwell M., 2011). Fungi that use plant tissues for reproduction and dispersal are among the most devastating plant pathogens, and over 19,000 fungi are characterized by great diversity and variability, which reduce crop yield and contaminate them with mycotoxins that harm human and animal health. *Fusarium spp*. are some of the most economically important fungal phytopathogens (Lazarovits G. et al., 2014). (Doehlemann G. et al., 2017; Möller M., Stukenbrock E.H., 2017).

Plant diseases caused by *Fusarium* species are mainly transmitted through soil and are difficult to control. One species of this genus, *Fusarium oxysporum*, is one of the top ten fungal plant pathogens that cause fungal infections in plants. It affects the yield, quality, and storage life of harvested plants. The significant economic impact of diseases caused by *Fusarium* determines the relevance of using fungicides in agriculture (Arie T., 2019; Dean R. et al., 2012; McGrath M.T., 2016; Steinberg G., Gurr S.J., 2020). However, along with the limited number of existing fungicides, their use in agriculture is limited due to toxicity and cross-resistance to medical antifungal agents (Lucas J.A. et al., 2015; Matsuzaki Y. et al., 2020; Brauer V.S. et al., 2019; Bastos R. W. et al., 2021). Therefore, there is a need for novel antifungal agents with high efficiency to control agricultural fungal diseases. Here we describe the activity of M451, which is a complex of the novel antimicrobial agent TGV-28 poly-N-carboxamido-1,6-diaminohexane) particularly N’-aminated and 0.01% NaH_2_PO_4_. Similar to its predecessor, Mul-1867, the mechanism of action of TGV-28 is a nonspecific attack on the cell walls of fungi (Tetz G., Tetz V., 2015; Tetz, G. et al., 2017). This study aimed to determine the in vitro activity of M451 against various phytopathogenic fungi.

## 2. Materials and Methods

### 2.1 Fungal isolates

The antifungal activity was tested against the following plant pathogenic fungi: *F. oxysporum, F. culmorum, F. graminearum, F. sporotrichioides, F. solani, F. verticillioides* (all from HMI collection, NY, USA), *F. proliferatum* KSU 4853 (Kansas State University, KS, USA), *F. dimerum* KSU 14971 (Kansas State University KS, USA), *Blumeria graminis, Claviceps purpurea, Alternaria alternata*, and *Phytophtora infestans* (all HMI collection).

### 2.2 Tested antifungals

The following antifungals were tested: Fludioxonil, amphotericin B, posaconazole, voriconazole, itraconazole, NaH_2_PO_4_ (Sigma–Aldrich, St. Louis, MO, USA), Maxim^®^ XL (Syngenta Canada Inc., Canada), M451 (Human Microbiology Institute, NY USA), which is a complex of T28 and 0.01% NaH_2_PO_4_ (Figure 1).

**Figure 1.**
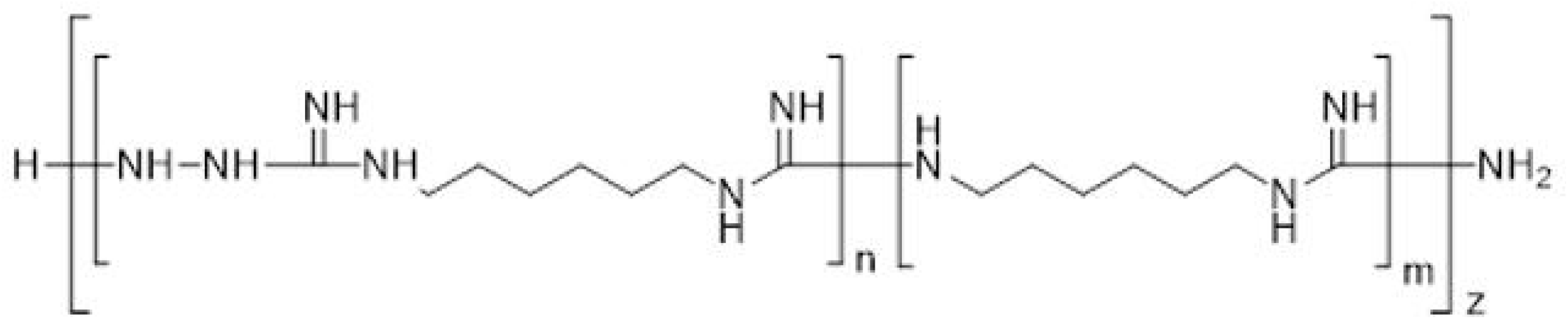
Chemical structure of TGV-28.

### 2.3 Growth media

Potato dextrose agar (PDA) (CM0139) (Thermo Fisher Scientific Inc., USA) and RPMI 1640 (Thermo Fisher Scientific, USA) were used.

### 2.4 *In Vitro* tests of mycelial growth inhibition

Mycelial growth inhibition was evaluated following the methodology reported by Chen F. et al. (2014) and Baggio J.S. et al. (2014). Briefly, varying concentrations of fungicidal agents were added to 90 mm Petri dishes filled with PDA (cooled to approximately 50 °C) before it solidified to obtain M451 0.1%, 0.05%, NaH_2_PO_4_ 0.1%, or fludioxonil 0.25%. The control plate contained untreated PDA medium. To test 13 fungal strains, mycelium plugs 3 mm in diameter of each fungal strain were cut from the edge of a 5-day-old colony grown on stock PDA plates, transferred to the control and experimental Petri dishes, and incubated at 25°C for 120 h. The diameter of the colonies was measured, and the percentage of growth inhibition was calculated.

### 2.5 Minimal fungicidal concentration (MFC) determination

The MFC of the tested products was assessed using the standard broth susceptibility testing method, according to the recommendations of the Clinical and Laboratory Standards Institute (CLSI) (Wayne P.A., 2008). The standard inoculum for fungal testing was taken as a fungal suspension of McFarland (0.5). MFC was defined as the lowest concentration of the tested compound, resulting in no visual growth after 48 h of incubation at 25°C. After 48 h of incubation at 25°C, the microdilution plates were examined for visible *Fusarium spp*. growth.

### 2.6 Kinetics of antifungal Activity

To determine the minimum concentration and exposure time required for the tested compounds to kill fungi, we performed a time-kill test in vitro. We assessed the activity of serially diluted compounds against *F. oxysporum* as described by Ernst et al., 1998. All fungal cultures were grown for 120 h as liquid cultures, and 150 μL of the inoculum (5 × 10^5^ CFU/mL) for each strain was transferred to 96-well microtiter plates (Sarstedt, Numbrecht, Germany). The plates were then incubated for 120 h at 25°C. Next, 50 μL of the tested compounds at different concentrations were added for 5 min, 30 min, 1 h, 3 h, 6 h, and 24 h. Untreated probes were used as negative controls at each time point. After the exposure, the fungi were centrifuged at 4000 × g for 15 min, washed with deionized water, and two additional rounds of centrifugation and washing were performed. The probes were diluted in 11 mL PBS, and the total CFU number was determined by serial dilution and plating on PDA. All assays included at least two replicates and were repeated in three independent experiments.

### 2.7 Spore germination assay

Spore germination was analyzed using a previously described method (Benslim A. et al. 2017), in which 1 mL of spore suspension (10^7^ spores/mL) was placed in a series of microtubes, and 20 µL of M451 was added. The tubes were prepared in triplicate and incubated for 24 h at 25 °C. After incubation, inhibition of spore germination was observed under an AxioStar Plus light microscope (Carl Zeiss, Germany; objective lens: A-Plan 100х/1.25) using a Malassez cell counting chamber (Thermo Fisher Scientific, USA). The percentage of non-germinated spores was calculated using the following equation:

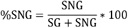

### 2.8 Spore production

Spore production was determined according to the procedure described by by Baggio J.S. et al., 2014. To test spore production by *F. oxysporum*, mycelial plugs 3 mm in diameter were taken from the edge of a 5-day-old colony and transferred into control and test tubes. The control tubes contained 10 mL of dH_2_0, 10 μL of Tween 20, and 30 μL of lactoglycerol as spore germination preventers. The test tubes contained the same mixture of fungicides at various concentrations. To count the number of spores using the Malassez counting chamber, we transferred 0.2 mL from each tube.

### 2.9 Germ tube elongation assay

The germ tube elongation was studied according to a previously reported procedure (Chen F. et al. 2014) To test the inhibition of germ tube elongation, we prepared a spore suspension of *F. oxysporum* (10^7^ spores/mL). One hundred microliters of this suspension were placed on control and experimental plates supplemented with M451 at various concentrations. The plates were incubated for 10 h at room temperature. Germ tube length was measured for 100 germinated spores at 400× magnification (Axiostar plus (Carl Zeiss, Germany) using Fiji software (Schindelin, J. et al., 2012).

### 2.10 Conidial destruction assay

The conidial destruction assay was evaluated following the methodology reported by Oukhouia M. et al. (2017). M451 (measured for TGV-28) and fludioxonil were diluted in 0.9% NaCl, mixed with 0.2% agar and added to 5-day-old *F. oxysporum* conidia suspension (10^6^ conidia/mL), up to a final concentration of 5000, 500, 100, 50, 5, 1, or 0.5 μg/mL. After 1 h, 3 h, 6 h, and 24 h of incubation at 25°C, the number of conidia was determined using a Malassez counting chamber. For the quantitative analysis, we used the following formula:

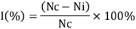

Conidia were counted in 10 different fields utilizing standard techniques (Niemann G.J. and Baayen R.P. 1989).

### 2.11 Statistical analyses

All data were analyzed using GraphPad Prism version 9.3.1. One-way analysis of variance (ANOVA) at a confidence level of p ≤ 0.05 was performed to determine the significance of the difference among treatments and concentrations, followed by Tukey’s multiple comparison test for the mean comparison test. All experiments were performed in triplicate.

## 3. Results

### 3.1 Antifungal activity of M451

First, we tested TGV-28 and NaH_2_PO_4_, alone and in combination with each other (M451), against different species from the phyla *Ascomycota, Oomycota*, and *Basidiomycota*. TGV28 was highly effective against all strains with MFC, varying from 64.3 to 96.3 µg/mL (Table 1). As expected, NaH_2_PO_4_ at 100 µg/mL did not exhibit antifungal activity. Surprisingly, the combination of TGV28 and NaH_2_PO_4_ increased the antifungal activity of TGV28 against *F. oxysporum* by 49% (p<0.05). Therefore, in subsequent experiments, we used a complex of TGV28 and NaH_2_PO_4_, named M451.

**Table 1.** Antifungal activity of tested compounds

| Fungi | Minimal Fungicidal Concentration (µg/mL) |  |  |
| --- | --- | --- | --- |
|  | TGV 28, µg/mL | NaH<sub>2</sub>PO<sub>4</sub>, | M451, µg/mL |

| | | $\mu\text{g/mL}$ | |
| --- | --- | --- | --- |
| *F. culmorum* | 1.3±0,6 | >1024 | 1.3±0,6 |
| *F. graminearum* | 0.2±0,3 | >1024 | 0.2±0,3 |
| *F. sporotrichioides* | 0.3±0,3 | >1024 | 0.3±0,3 |
| *F. oxysporum* | 5.3±2,3 | >1024 | 2.7±1,2 |
| *F. solani* | 2.7±1,2 | >1024 | 2.7±1,2 |
| *F. dimerum* | 3.3±1,2 | >1024 | 3.3±1,2 |
| *F. proliferatum* | 1.3±0,6 | >1024 | 1.3±0,6 |
| *F. verticillioides* | 0.8±0,3 | >1024 | 0.8±0,3 |
| *Erysiphe graminis* | 1.3±0,6 | >1024 | 1.3±0,6 |
| *Claviceps purpurea* | 0.2±0,3 | >1024 | 1.3±0,6 |
| *Alternaria alternata* | 1.7±0,6 | >1024 | 1.7±0,6 |
| *Phytophthora infestans* | 0.3±0,3 | >1024 | 0.2±0,3 |
| *Rhizoctonia solani* | 0.3±0,3 | >1024 | 0.3±0,3 |

### 3.2 Mycelial growth inhibition assay

We found that M451 0.1% had the highest fungicidal activity against all tested 13 fungal strains of the *Ascomycota, Oomycota*, and *Basidiomycota* phyla, displaying 92.6–95.4% mycelial growth inhibition (all p<0.05) (Figure 2A, Supplementary Table 1). A slightly smaller decrease in mycelial growth was noted when M451 0.05% was used, with 82.7 to 85.9% inhibition across different fungal pathogens. Notably, both M451 0.05% and 0.1% were more effective against any of the tested strains than fludioxonil (the active ingredient in Maxim XL) at a fixed 0.25% concentration according to the manufacturer’s instructions (Figure 2A–E).

**Figure 2.**
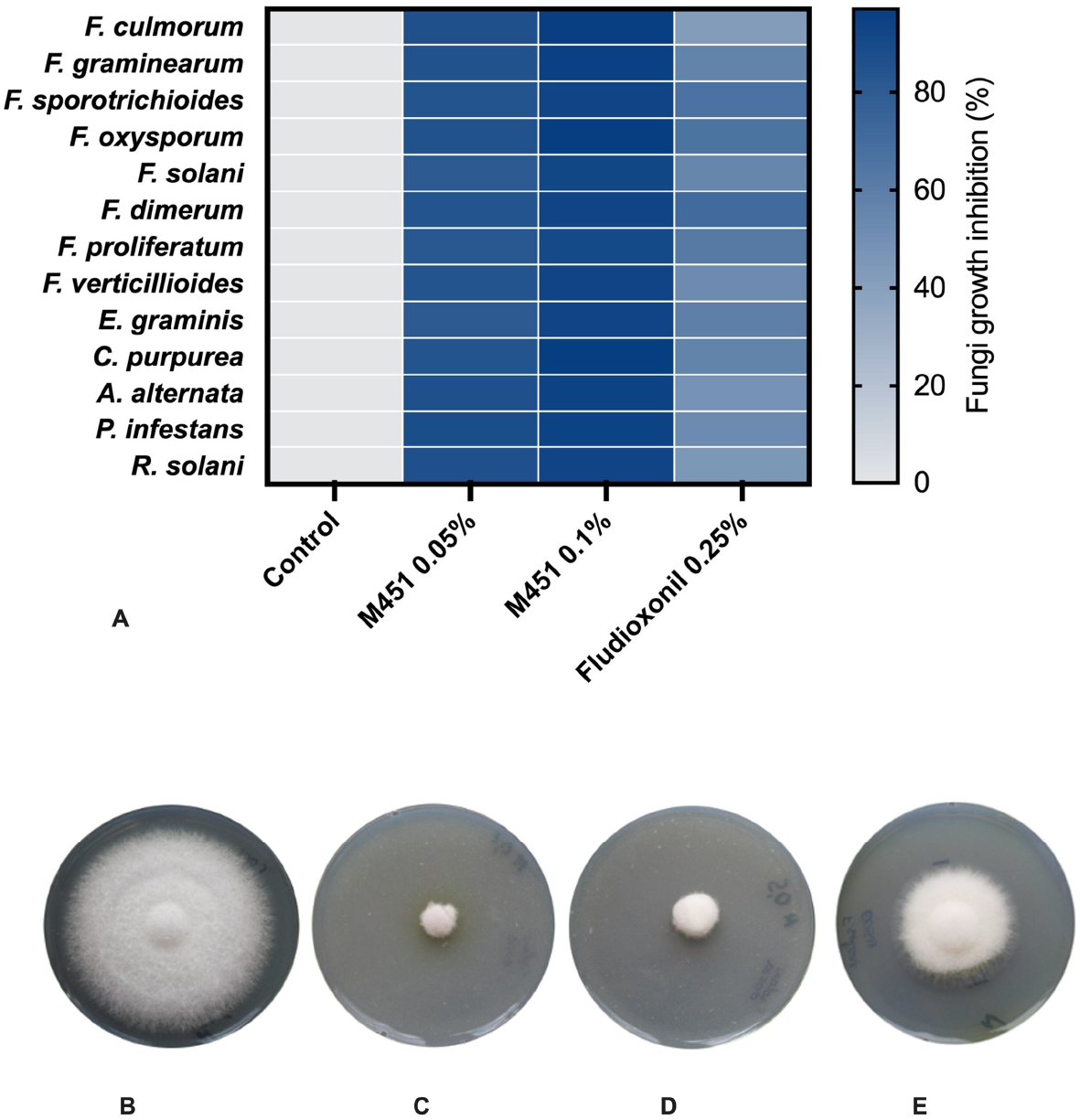
Effect of tested compounds on fungal growth (**A**). Mycelial growth inhibition of 13 tested fungal strains of *Ascomycota, Oomycota* and *Basidiomycota* phylum. (**B–E**) representative images of antifungal activity of M451 against *F. oxysporum*. Growth 5 days at 25 °C after the treatment with M451 **(B)** negative control **(C)** M451 0.1% **(D)** M451 0.05% **(E)** Fludioxonil 0.25%. The figures are representative images from three independent experiments.

### 3.2 The median effective concentration (EC50) value of M451 against different plant pathogens

M451 exhibited significant activity against all tested plant-pathogenic fungal isolates. We found that it inhibited the in vitro mycelial growth of all strains of *Ascomycota, Oomycota*, and *Basidiomycota* (Table 2), with an EC_50_ of 34–145 µg/mL and a high correlation coefficient. The EC_50_ of M451 against *Fusarium spp*. (phylum *Ascomycota*) varied from 66 µg/mL to 145 µg/mL. Representative images of the mycelial growth inhibition of *F. oxysporum* by M451 are shown in Figure 1. The activity against other non-*Fusarium* representatives of the phylum *Ascomycota* was higher, with EC50 values ranging from 34 to 52 µg/mL. The EC50 values for *Phytophtora infestans* (phylum *Oomycota*) and *Rhizoctonia solani* (phylum *Basidiomycota*) were 58 and 53 g/mL, respectively.

**Table 2.**
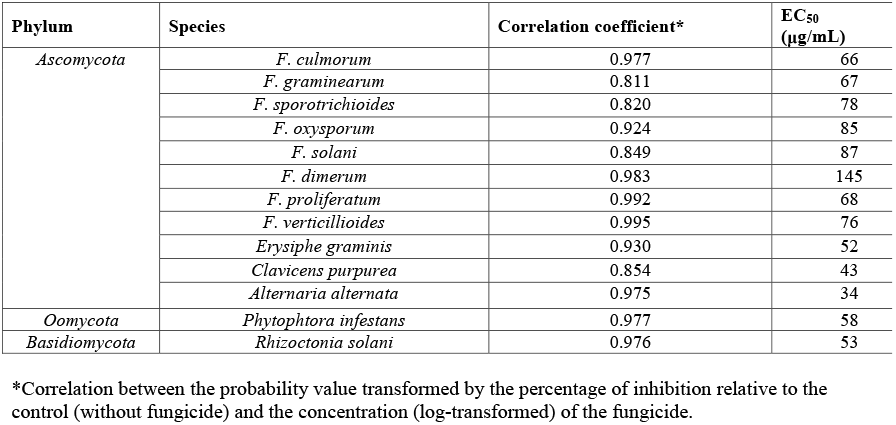
In vitro inhibition of plant pathogens by M451

### 3.3 The MFC of M451 and different antibiotics against *F. oxysporum*

Table 3 shows the MFCs of M451 and antifungal antibiotics representing polyene, azole classes, and stand-alone Maxim XL (titrated depending on the fludioxonil concentration) against *F. oxysporum*. The results of the time-kill study against planktonic fungi demonstrated higher antifungal activity of M451 at all tested time points. At the end of 24 h of exposure, the antifungal activity of M451 was 5–50 times higher than that of other antifungal agents.

**Table 3.** The MFCs of M451 and antifungal antibiotics against *F. oxysporum*

| Antifungal agent | MFC at different incubation ($\mu\text{g/mL}$) | | | | | |
| --- | --- | --- | --- | --- | --- | --- |
|  | 5 min | 30 min | 1h | 3h | 6h | 24h |
| M451 | 512 | 64 | 32 | 16 | 16 | 4 |
| Fludioxonil | >1024 | >1024 | 512 | 256 | 256 | 256 |
| Amphotericin B | >1024 | >1024 | 256 | 128 | 64 | 32 |
| Posaconazole | >1024 | >1024 | 128 | 64 | 32 | 32 |
| Voriconazole | >1024 | >1024 | 128 | 128 | 128 | 32 |
| Itraconazole | >1024 | >1024 | 256 | 128 | 64 | 32 |

### 3.4 Conidial destruction assays

Next, we studied the effects of different concentrations of M451 corresponding to 5000–5 µg/mL TGV-28 compared with Maxim XL (titrated according to fludioxonil concentration) on *F. oxysporum* conidia in conidial destruction assays (Figure 3, Supplementary Table 2). M451 treatment at 5000 µg/mL and 500 µg/mL induced growth inhibition by 99% and 97.8%, respectively, after 5 min of contact time. We could clearly see the dose- and time-dependent effects of M451 at lower concentrations. Thirty minutes of incubation with M451 at 100 µg/mL resulted in 53.8% inhibition; increasing the incubation time to three hours resulted in 94.4% inhibition. Under the same conditions, Maxim XL had a much lower level of anti-*Fusarium* activity. Even at a 5000 µg/mL concentration, it only reduced viability by 3.6% after 5 min and 67.8% after 24 h.

**Figure 3.**
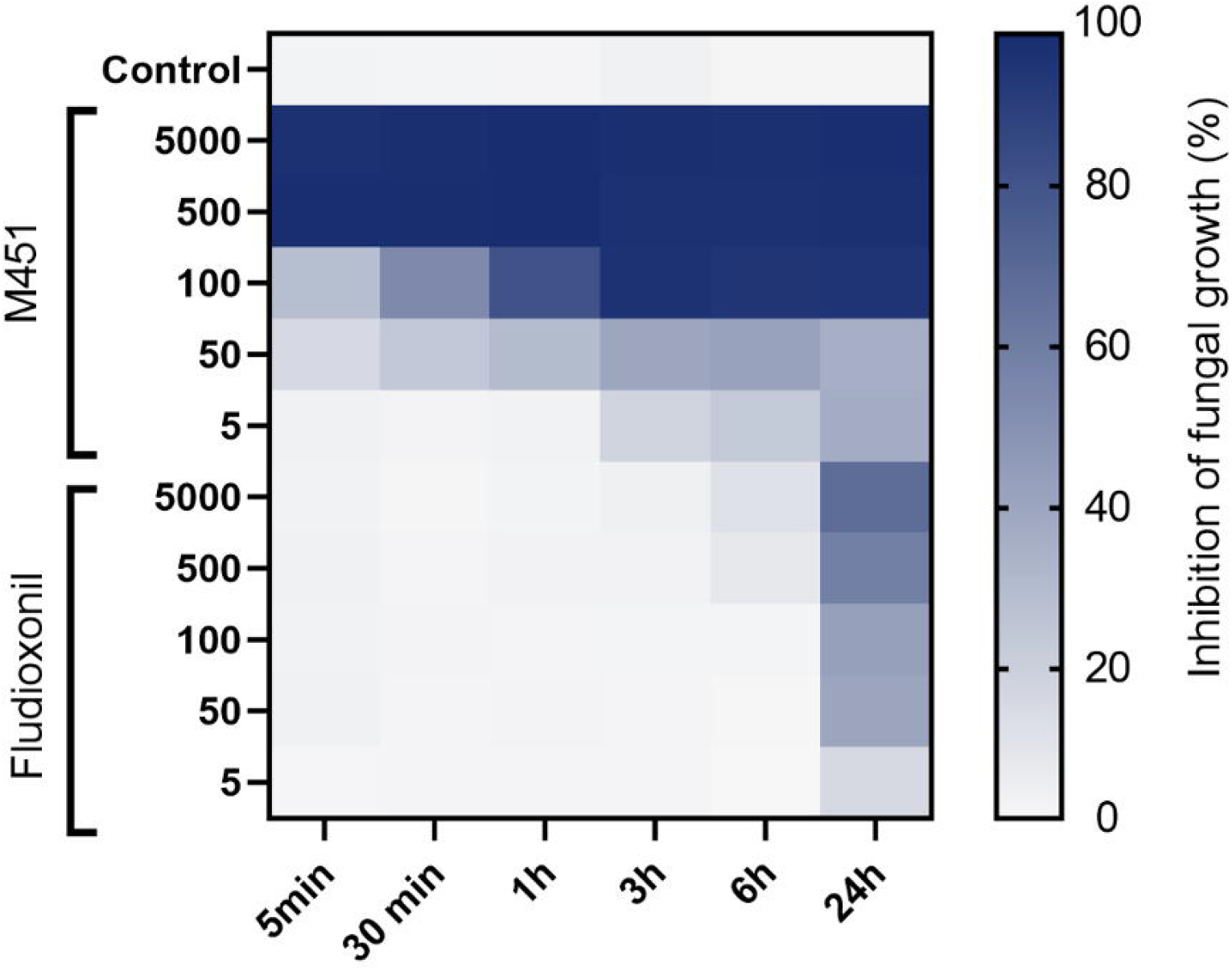
Heatmap of the effect of different concentrations of M451 and fludioxonil on *F. oxysporum* conidia. Growth inhibition was represented by a color scale from white (minimal) to blue (maximum). Data are representative of three independent experiments.

### 3.5 Effect of M451 on different developmental stages of *F. oxysporum*

Next, we determined the in vitro effects of M451 on germ tube elongation, spore production, and germination of *F. oxysporum*. (Figure 4). Germ tube elongation was the most sensitive to M451, with over 80% inhibition, even when M451 was applied at a sub-inhibitory concentration of 50 µg/mL (Figure 4A). Notably, M451 significantly inhibited spore production at all the tested concentrations. This might be explained by the quick action of M451 when the compound kills fungi faster than the sporulation process starts (Figure 4B). In addition, we observed that M451 had a dose-dependent inhibitory effect on *Fusarium* spore germination (Figure 4C).

**Figure 4.**
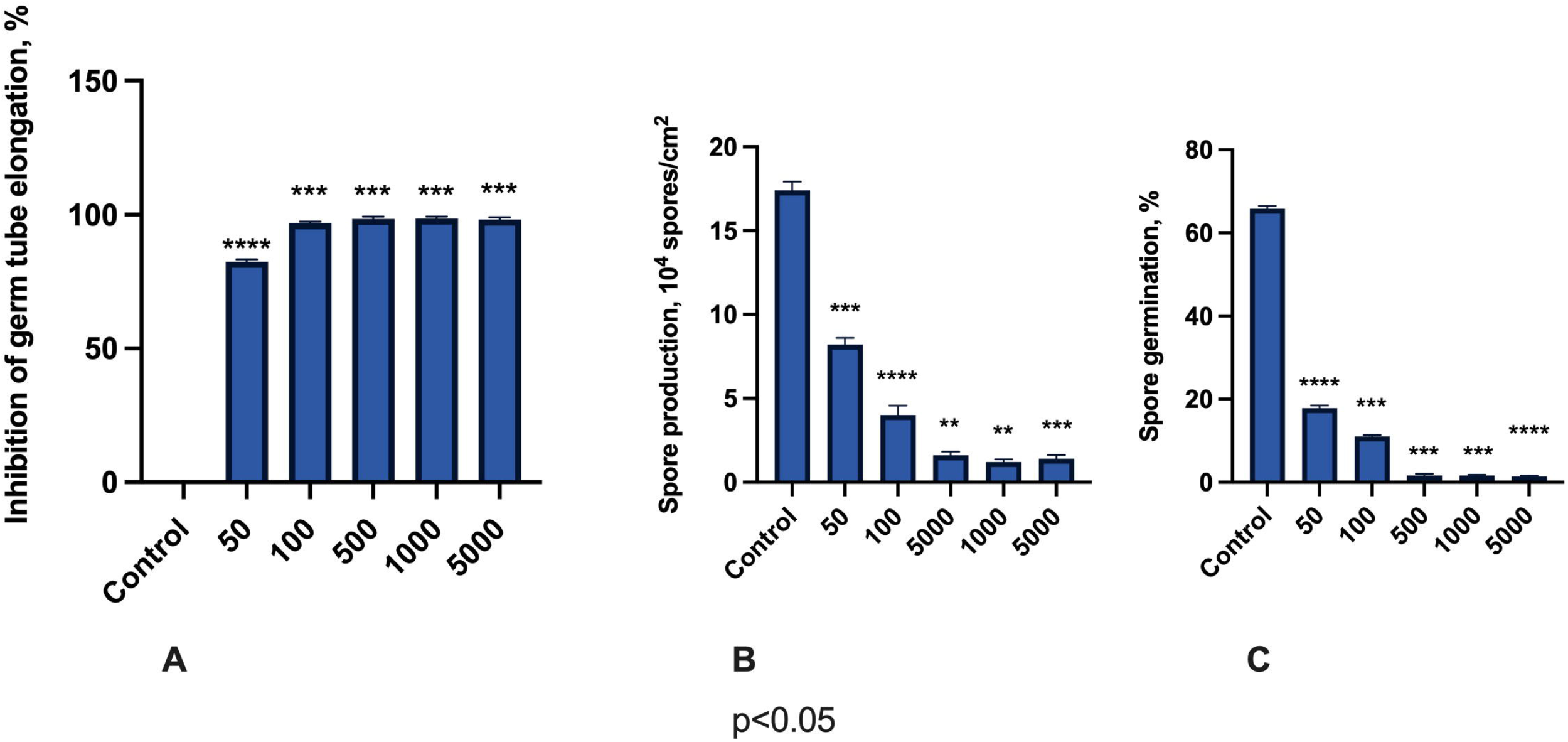
Antifungal effects of M451 on *F. oxysporum*. The bars represent the effects of M541 at concentrations from 50 to 5000 µg/mL on **(A)** germ tube elongation, **(B)** spore production, and **(C)** spore germination. **p<0.01, ***p<0.001, ****p<0.0001. Compiled data are from three independent experiments.

## 4 DISCUSSION

*Ascomycota, Oomycota*, and *Basidiomycota* spp. cause many diseases with an economic impact on important agronomic host plants and represent a significant challenge for agriculture (Haverkort A.J. et al., 2016; Tudzynski P. et al., 2003; Gisi U., Cohen Y., 1996). The main reasons for the difficulties are the pathogen’s resistance to the action of existing fungicides and insufficiently studied regulations (Tetz G., Tetz V., 2022; Tetz G., Tetz V., 2022; Jindo K., 2021; Al-Hatmi A.M. et al., 2019Meyers E. et al., 2019; Avenot H.F., Michailides T.J., 2007; Gisi U., Cohen Y., 1996; Bennett R.S. et al., 2011).

In this study, we found that the novel fungicide candidate, TGV28, exhibits a high level of antifungal activity against many emergent plant pathogens of the phyla *Ascomycota, Oomycota*, and *Basidiomycota*. However, in this study, we used a complex of TGV28 and NaH_2_PO_4_, named M451. Although NaH_2_PO_4_ had no antifungal activity on its own, it enhanced the antimicrobial activity of TGV-28 against certain fungal pathogens. The mechanisms behind this action are a subject for future research.

In time-kill studies, when we compared the activity of M451 with different antifungal agents, including azole derivatives and fludioxonil (an antifungal component of Maxim XL) and found that M451 had both a higher level and quicker-acting level of antifungal effects. This is because TGV-28 exhibits a topical mechanism of antifungal action (Tetz G et al.,2017). Therefore, M451-induced rapid killing of fungal cells and prevented their survival through sporulation. However, under the same conditions in the conidial destruction assay, fludioxonil only had antifungal activity after 24 h of treatment, even at the highest concentration. This indicates that fludioxonil failed to prevent sporulation.

Moreover, M451 also inhibited the germination of *F. oxysporum* spores. Previous studies have revealed that this class of compounds does not have significant cytotoxicity; therefore, the observed sporicidal effect could not be addressed due to corrosive effects (Tetz G et al., 2017). The high efficacy of M451 against strains of difficult-to-treat plant pathogens, along with its lack of cross-resistance and high sporicidal activity, means that this compound may be employed as an agricultural fungicide for the successful control of fungal diseases in plants.

## 5 Conclusion

In this study, we investigated the antifungal activity of the novel antifungal agent, M451, against different plant pathogens. M451 substantially inhibited mycelial growth, germ tube elongation, sporulation, and spore germination, with the antifungal activity exceeding that of the azole derivatives of fludioxonil. Also, our results indicated that M451 had a rapid effect and inhibited more than 97% of the mycelial growth of *F. oxysporum* within 5 min. The inhibitory effect of M451 on the development of different plant pathogens indicates its potential as a novel chemical fungicide.

## 6 Data availability statement

The original contributions of this study are included in this article. Further inquiries can be directed to the corresponding authors.

## 8 Author Contributions

VT, KKM, GT conceived and supervised the research. KKM and MV conducted experiments. KKM and KK analyzed the data and wrote the manuscript. KKM, GT, VT edited and helped to draft the final manuscript. All authors contributed to the article and approved the submitted version.

## 10 Acknowledgments

Nothing to declare.

